# Sulfisoxazole does not inhibit the secretion of small extracellular vesicles

**DOI:** 10.1101/2020.03.15.988055

**Authors:** Pamali Fonseka, Sai V Chitti, Rahul Sanwlani, Suresh Mathivanan

**Affiliations:** Department of Biochemistry and Genetics, La Trobe Institute for Molecular Science, La Trobe University, Melbourne, Victoria, 3086, Australia

**Author notes:** To whom correspondence should be addressed Professor Suresh Mathivanan, Department of Biochemistry and genetics, La Trobe institute for molecular Science, La Trobe University, Bundoora, Victoria 3086, Australia. Authors contributed equally (Pamali Fonseka, Sai V Chitti, Rahul Sanwlani).

## Abstract

Recently, the study by Im et *al*. focused on blocking the release of extracellular vesicles (EVs) by cancer cells, as a strategy to block metastasis, by deploying a drug repurposing screen. Upon screening the library of FDA approved drugs in breast cancer cells *in vitro*, the authors reported the ability of the antibiotic Sulfisoxazole (SFX) in inhibiting EV biogenesis and secretion. SFX was also effective in reducing breast primary tumor burden and blocking metastasis in immunocompromised and immunocompetent mouse models. As we seek a compound to block EV biogenesis and secretion in our current *in vivo* studies, we intended to use SFX and hence performed *in vitro* characterization as the first step. However, treatment of two cancer cells with SFX did not reduce the amount of EVs as reported by the authors.

---

Over the last two decades, extracellular vesicles (EVs) have been implicated in intercellular communication and utilised as drug delivery vehicles and as reservoirs of disease biomarkers^1-5^. As they continue to garner interest, several seminal studies have established that EVs regulate various pathophysiological processes in favour of cancer progression, including remodelling the tumour microenvironment, immune evasion, coagulation, vascular leakiness, establishing the pre-metastatic niche, tropism for metastasis and transfer of chemoresistance^6-11^. Hence, there is growing interest in blocking the release of EVs and limiting their systemic circulation as a novel therapeutic avenue to treat cancer^12^. As anti-metastatic therapies are scarce, it is speculated that FDA approved drugs that target EVs could possibly fill the void.

Recently, the study by Im *et al*. focused on blocking the release of EVs by cancer cells, as a strategy to block metastasis, by deploying a drug repurposing screen^13^. The rationale was to screen the existing FDA approved library to identify drugs which can inhibit EV biogenesis or secretion with the obvious advantage of known mode of action, efficacy and toxicity profile and hence has the potential of immediate clinical utility. Upon screening the library of FDA approved drugs in metastatic breast cancer cells *in vitro*, the authors reported the ability of the antibiotic Sulfisoxazole (SFX) in inhibiting EV biogenesis and secretion. The authors also reported that SFX was effective in reducing breast primary tumor burden and blocking metastasis in immunocompromised and immunocompetent mouse models. SFX was proposed to target Endothelin receptor A (ETA) previously in 1994^14^ and hence the authors validated that SFX targets ETA which can positively regulate EV biogenies and secretion. The findings in this study thus present SFX as a potential novel EV-targeted therapeutic alternative. As a group interested in EVs, the outcomes proposed in the study were encouraging and attractive as FDA approved drugs targeting EV release are limited. Recently, Datta *et al*. also performed a repurposing screen to identify drugs that modulate the release of EVs in prostate cancer cells^12^. However, the identified drugs are yet to be tested *in vivo*.

As we seek a compound to block EV biogenesis and secretion in our current *in vivo* studies, we intended to use SFX and hence performed *in vitro* characterization as the first step. However, treatment of 4T1 breast cancer cells with SFX did not reduce the amount of EVs as reported by the authors. We acknowledge the fact that our EV isolation protocol^15^ was different from the study^13^ (**Fig. 1** – Mathivanan laboratory protocol) and hence could have attributed to the varied results. In order to rule out the possibility of variations in the method of EV isolation or cell-type dependency, three researchers exactly followed the protocol employed by the authors (**Fig. 1** – protocol 1 and 2) to isolate EVs from two different cell types (4T1 and MDA-MB231) that were used by the authors. All these assays were performed with 3 technical replicates (same batch of cells seeded in three different plates) for every biological replicate. Consistent with our previous observations, treatment of the cancer cells with varying concentrations of SFX (50, 100 and 200 µM) did not impede the release of EVs while the positive control ceramide inhibitor GW4869, at low concentration (5 µM), inhibited EV secretion. Upon EV isolation, we quantified the protein amount, particle number and performed Western blotting for EV enriched proteins (TSG101, Alix), all normalised to equal cell number (**Fig. 2**). Contrary to the authors claim of SFX treatment led to a 3-fold decline in EV particle number, we observed a significant increase in particle number upon SFX (200 µM) treatment.

**Figure 1.**
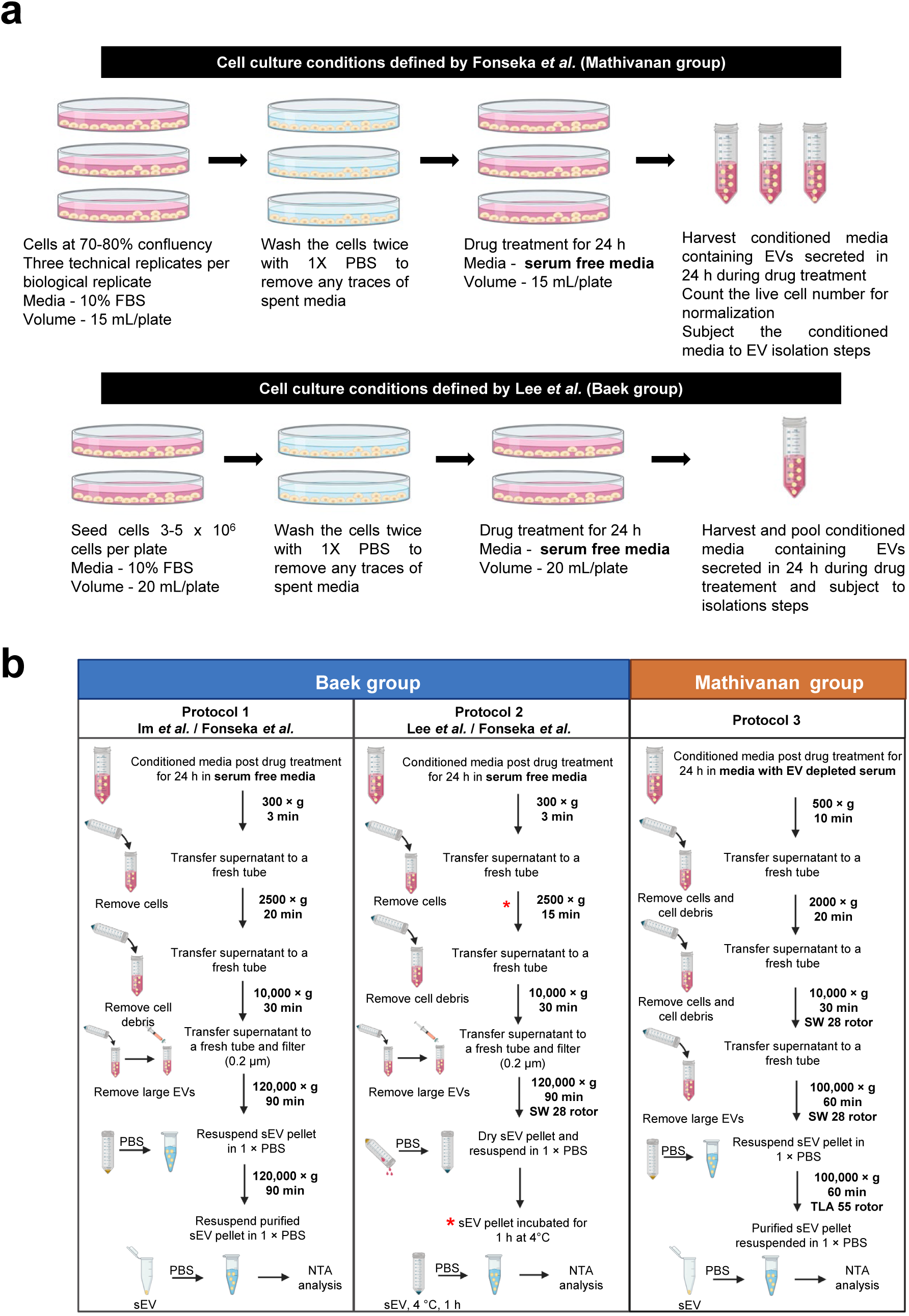
Schematic representation of cell culture, drug treatment and sEV isolation by two groups. **(a)** Schematic representation of cell culture and drug treatment approaches highlighting our approach. The cells were seeded and allowed to grow to 70-80% confluency in media supplemented with FCS. Upon attaining desired confluency, cells were washed with 1 × PBS and subjected to drug treatment for 24 h in serum free media. At treatment endpoint, conditioned media was harvested and subjected to sEV isolation. Live cell number was determined to normalise particles released to cell number. Similarly, Baek group seeded cells in a pre-defined range and treated them with drug in serum free media. **(b)** Protocol 1, schematic flow diagram depicting the methodology of sEV isolation and analyses as defined by Baek group in the initial publication (Im et al.) and followed by Fonseka *et al*. Protocol 2, schematic flow diagram depicting the methodology of sEV isolation and analyses as defined by Baek group in the latest version (Lee et al.). The variations in the methodology from the original manuscript (Im et al.) is marked (*). Protocol 2 was also adapted by Fonseka *et al*. to include the variations (*) suggested by Baek group. Protocol 3, standardized and optimized method of sEV isolation in EV depleted serum used in Mathivanan laboratory was also used to collect sEVs post treatment with SFX.

**Figure 2.**
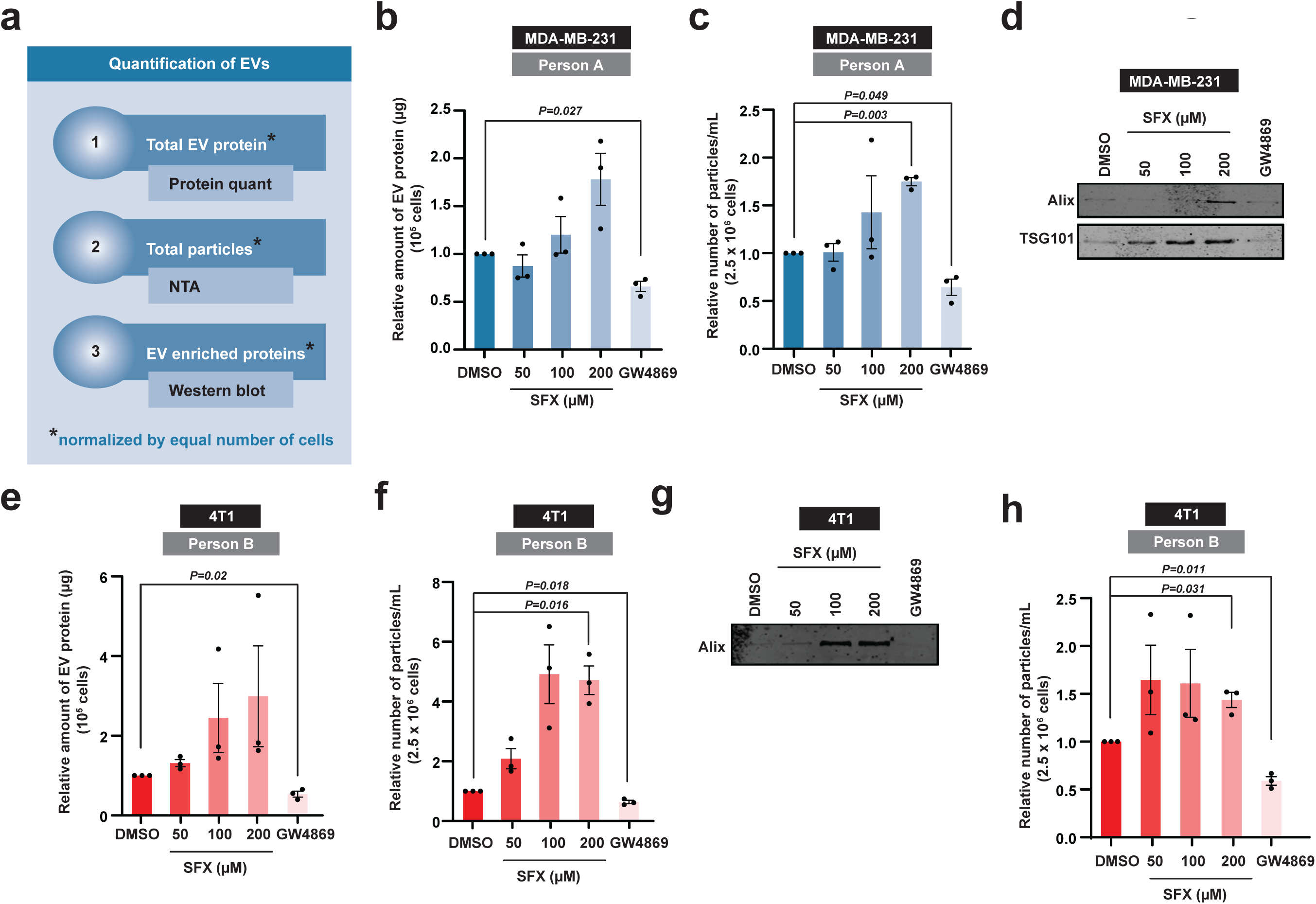
Quantification of EVs released by cells with or without SFX and GW4869. **(a)** Schematic of EV quantification by three different methods. Firstly, the total EV protein amount was quantified and normalised to equal number of live cells. Secondly, nanoparticle tracking analysis (NTA) was performed to quantify the total number of particles normalised to equal number of live cells. Lastly, Western blot analysis of EV samples obtained from equal number of live cells was performed for EV enriched proteins. **(b)** Relative amount of EV protein normalised to 10^5^ MDA-MB-231 cells is shown. **(c)** Relative number of particles normalised to 2.5 × 10^6^ MDA-MB-231 cells is depicted. **(d)** Western blot analysis of EV enriched proteins Alix and TSG101 in EV samples obtained from 15 × 10^6^ MDA-MB-231 cells. Representative image shown, *n=3*. **(e)** Relative amount of EV protein normalised to 10^5^ 4T1 cells is shown. **(f)** Relative number of particles normalised to 2.5 × 10^6^ 4T1 cells is depicted. **(g)** Western blot analysis of EV enriched proteins Alix in EV samples obtained from 15 × 10^6^ 4T1 cells. Representative image shown, *n=3*. **(h)** Relative number of particles normalised to 2.5 × 10^6^ 4T1 cells is depicted. All data are represented as mean ± s.e.m. *n=3*, statistical significance was determined by paired two-tailed t-test. EVs were isolated by protocol 1 (**b-g**) and 2 (**h**).

Overall, we report that SFX does not reduce the release of EVs (3 independent researchers) and emphasise caution in using SFX (Merk 31739) as a drug to block EV release. However, we do acknowledge the fact that our findings do not challenge the authors main conclusion of SFX mediated reduction of primary tumour burden and metastasis though our results suggest that the phenotype observed may not be cancer cell-derived EV mediated. Further research is needed to understand as how SFX can reduce primary tumor burden and inhibit metastasis.

## Acknowledgements

Suresh Mathivanan is supported by Australian Research Council Future Fellowship (FT180100333). The funders had no role in study design, data collection and analysis, decision to publish, or preparation of the manuscript. Fig. 1 has been created with Biorender.com.

## Author contributions

S.V.C. and S.M. conceived the concept. P.F., S.V.C. and R.S. performed the experiments. P.F. and S.M. made the figures. R.S. and S.M. wrote the manuscript. All authors read and approved the manuscript.

